# Personalized Tumor Growth Prediction with Multiscale Tumor Modeling

**DOI:** 10.1101/510172

**Authors:** Serbulent Unsal, Aybar Acar, Mehmet Itik, Ayse Kabatas, Oznur Gedikli, Feyyaz Ozdemir, Kemal Turhan

## Abstract

**Background:** Cancer is one of the most complex phenomena in biology and medicine. Extensive attempts have been made to work around this complexity. In this study, we try to take a selective approach; not modeling each particular facet in detail but rather only the pertinent and essential parts of the tumor system are simulated and followed by optimization, revealing specific traits. This leads us to a pellucid personalized model which is noteworthy as it closely approximates existing experimental results.

**Method:** For years, research has focused on modeling tumor growth but not many studies have put forward a framework for the personalization of models. In the present study, a hybrid modeling approach which consists of cellular automata for discrete cell state representation and diffusion equations to calculate distribution of relevant substances in the tumor micro-environment is favored. Moreover, naive Bayesian decision making with weighted stochastic equations and a Bayesian network to model the temporal order of mutations is presented. The model is personalized according to the evidence using Markov Chain Monte Carlo. Ultimately, this way of thinking about tumor modeling leads us to a vascular multi-scale model of tumor growth.

**Results:** To validate the tumor model, a data set belonging to the A549 cell line is used. The data represents the growth of a tumor for 30 days. We optimize the coefficients of the stochastic decision making equations using first half of the timeline. Then we predict next 15 days of growth without any other supervision. Results are promising with their low error margin and simulated growth data is in line with laboratory results.

**Conclusion:** There are many subsystems which have an effect in the growth of a tumor. A detailed model which includes all of them is currently virtually impossible to implement. We have therefore focused on a system that only includes fundamental components in this study, and have evaluated its predictions. We propose novel probability functions to obtain a personalized model and estimate the individual importance (weights) of each with parameter optimization. Our approach of using simulated annealing for parameter estimation and the subsequent validation of the prediction with in-vitro tumor growth data are, to our knowledge, unique in the literature.

## 1 Background

Despite much progress in oncology, molecular biology, and related fields, cancer is still a condition for which the prognosis is generally a shortened lifespan or lowered quality of life, frequently dramatically so. The complex and individually particular behavior of cancer decreases success rates of cancer therapies. The usual steps of cancer therapy are: deciding on tumor’s pathological type, staging the cancer using clinical data and planning the therapy according to medical guidelines which are informed by bulk statistics. In this routine, there is little room to calculate and predict patient’s therapy response in a bespoke way.

Mathematical models that use patient specific data and up-to-date scientific evidence has implications for the evidence based practice of personalized medicine. Use of individually tuned mathematical models give clinicians the ability to compare alternative therapy plans and predict outcomes. These models have an important role in the early drug development and for development of therapy scheduling. However, this search for personalized therapy has not yet met success. In the present study we propose a hybrid tumor model for the Non-Small Cell Lung Cancer (NSCLC) and a personalization framework.

Different approaches exist to develop tumor models. Continuous tumor models simulate tumor growth with in a set of differential equations, making them good options in modeling complex systems [1–6]. Simulating attributes of a tumor at tissue scale is trivial with continuous models. However, it is a non-trivial challenge to use them for simulating individual cell dynamics or discrete events in a cell or in the cell’s microenvironment. Discrete models are a solution to this problem. Simulation of the tumor system dynamics, which cannot be easily modeled by the continuous approach can be possible with the discrete modeling techniques such as the orientation mechanism of tumor according to nutrients [7], effects of the cell adhesion on tumor growth as well as effects of the proteolytic enzymes [8], competition between cell colonies [9], genetic parameters on the tumor movement [10]. Hence, it could be concluded that, discrete models provide sufficient flexibility in modeling cell-state scale tumor dynamics. Continuous and discrete modeling approaches could be thus combined into hybrid models [11]. Most of the time a discrete model is created at cell scale to simulate behavior of the cells and a continuous model is used at tissue scale to simulate distribution of substances like oxygen or glucose in tumor micro environment in hybrid modeling [9, 12].

In this study, such a framework has been developed for personalized tumor modeling that includes tumor and tissue specific parameters gathered from the literature for A549[13], which is a well known cell line derived of lung adenocarcinoma. Cellular automata are used for discrete cell state representation. Substance distribution in the vascular tumor micro-environment is calculated by using partial differential equations. Mutations of cells are modeled with a probabilistic network. A Naive Bayes approach is chosen for the decision making module of each cell. Weighted stochastic equations are created for modeling decisions of the tumor cells. Overall, this approach enables us to create a model which can easily be personalized by the optimization of individual parameters using simulated annealing. Results are promising with their low error margin and the simulated growth data fits well to xenograft model.

### State of The Art

According to the best of our knowledge;

- Our model is the first such model able to accurately regress the personalized growth of lung adenocarcinoma given data from the stages.
- Our approach of personalization uses simulated annealing and its validation with a xenograft model is novel.
- The model also incorporates a hierarchical Bayesian network of the tumor which is created from A549 mutation data and uses this model to predict the order of occurrence and timing of consecutive mutations during tumor progression. Although there are a few models which use mutation data, our model uses temporal and hierarchal order of specific cancer driver genes which has, thus far, to out knowledge, has not been leveraged.
- We propose novel probability functions to obtain a personalized model and estimate the importance (weight) of each with optimization.

## 2 Results

### 2.1 Consistency

A tumor model should be consistent with the basics of tumor biology. We first test this consistency using experimental data. Studies show that tumor growth accelerates under hypoxia [14–17]. Figure 1 shows our model is consistent with literature for hypoxic growth dynamics. Pearson correlation coefficient calculated between model and study for hypoxic and normoxic conditions. A strong correlation for both hypoxic (r=0.992) and normoxic (r=0.991) conditions is observed on tumor volume. Also as seen in Figure 2 in hypoxia which could be defined as 10% *O*_2_ of normoxia [18] apoptic region is considerably larger and tumor shows migrative behavior which is also in line with experimental results [19, 20]. In the figure,

**Figure 1.**
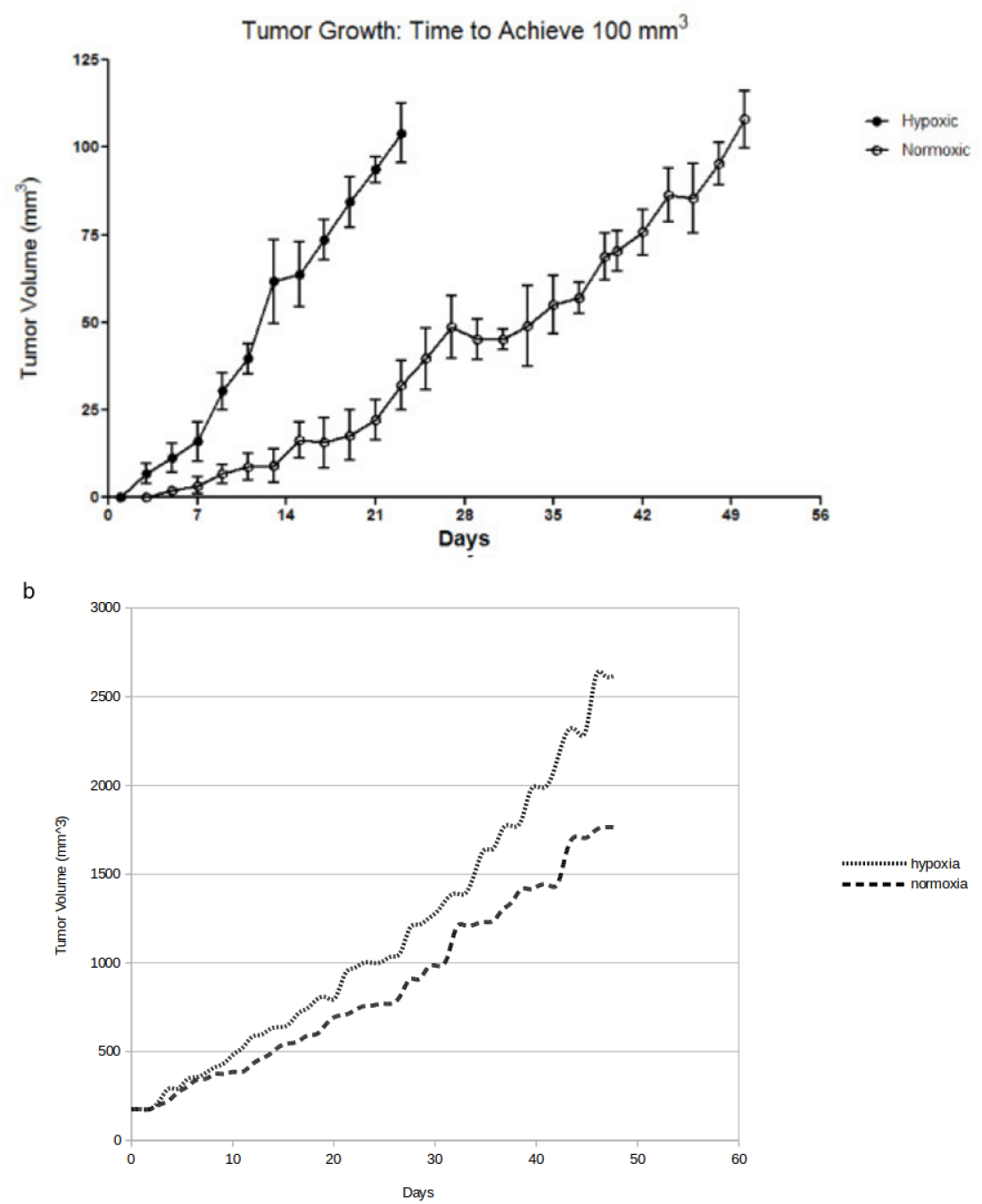
**a** [14] reports tumor growth under hypoxic conditions (10% *O*_2_) **b** Our *in-silico* experiment results

**Figure 2.**
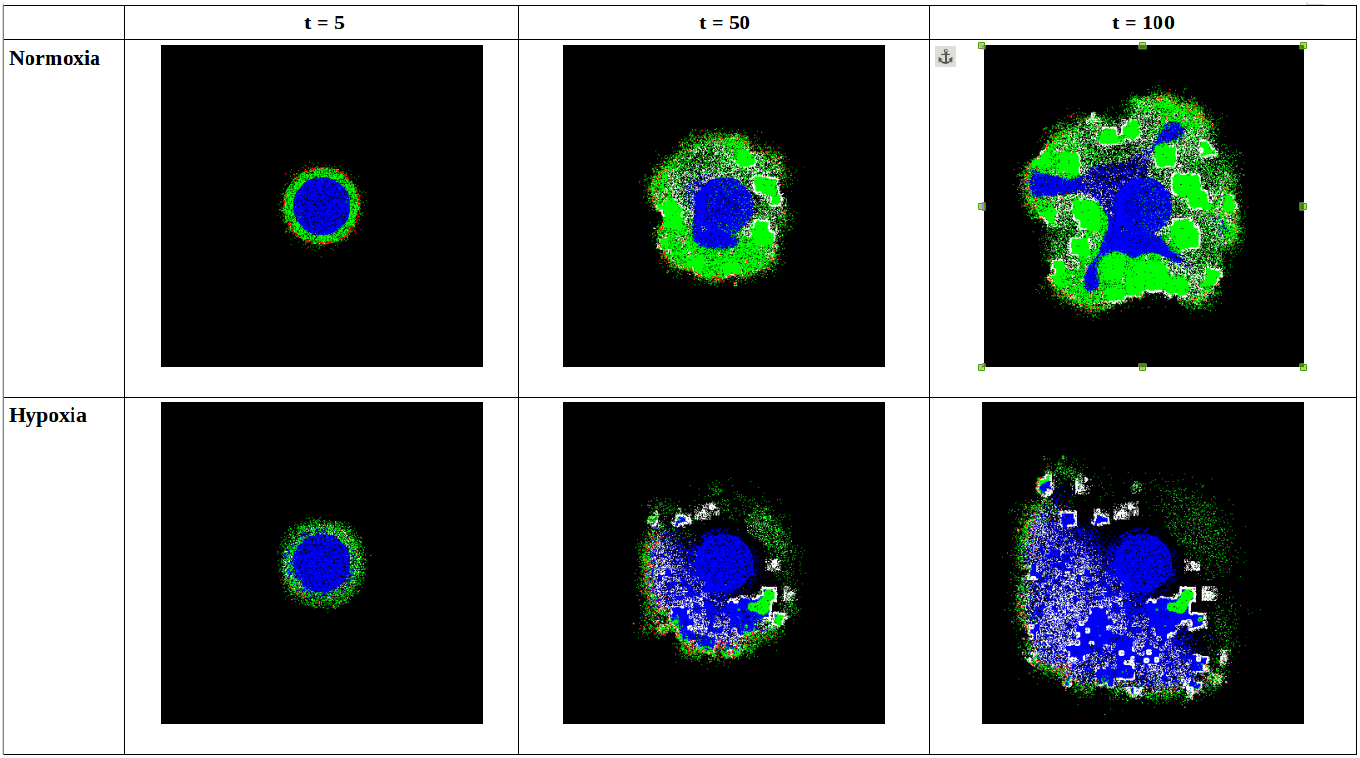
Tumor growth at time steps 5, 50 and 100 for hypoxia and normoxia. Time step t equals to 22 hours. Red cells have proliferative, green cells have quiescent, and white cells have migrative phenotypes while blue represents non-viable cells, whether apoptotic or necrotic.

Many studies have shown that glucose is an important factor in tumor growth. This fact was shown not only in-vitro [21–24]; but also in-vivo experiments [25]. Our model is in line with these studies as shown in Figure 3. Pearson correlation coefficient calculated between model and study for normal and low glucose conditions. A strong correlation for both normal (r=0.998) and low glucose (r=0.898) conditions is observed on tumor cell number. Limitation of growth under low glucose can also be observed by comparing tumor morphology as seen in Figure 4.

**Figure 3.**
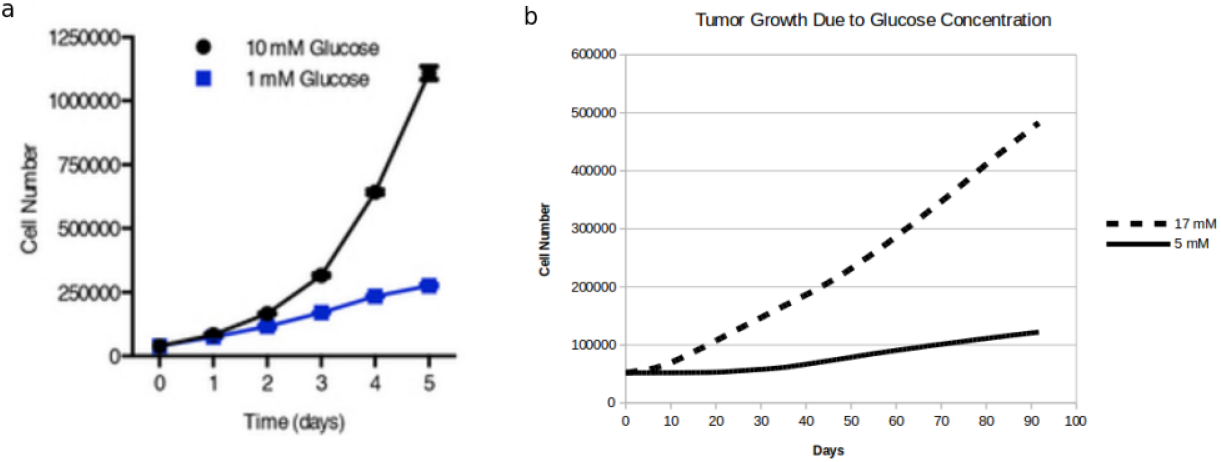
**a** Previously reported[14] tumor growth under different glucose concentrations. **b** Results of our *in-silico* results.

**Figure 4.**
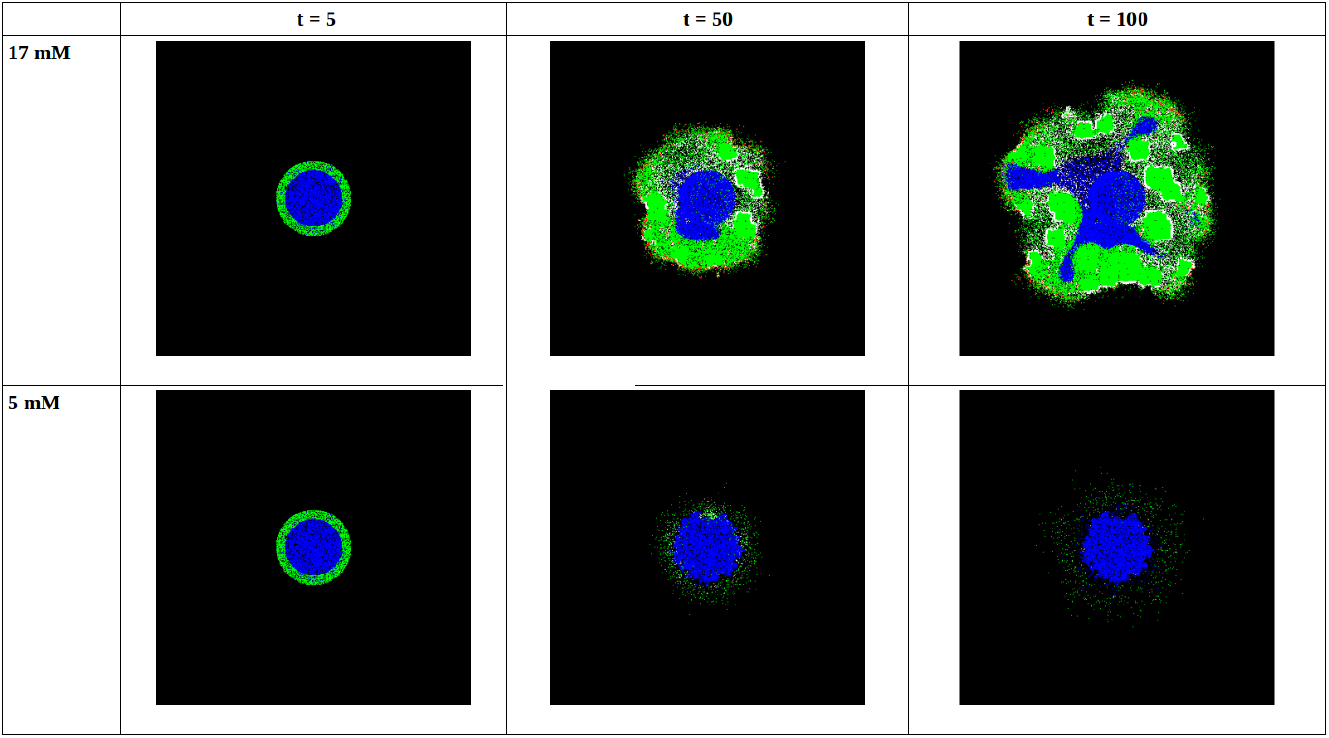
Tumor growth at time steps 5, 50 and 100 for normal (17 mM) and low (5 mM) glucose levels. Time step t equals to 22 hours.

Chemotaxis is a fundamental mechanism that determines tumor morphology: defined as motility of cells towards resources like oxygen and glucose. To ensure that cells in our model simulate chemotactic behavior we created a set of capillaries as shown in Figure 5. Subsequently, the growth of the simulated tumor was monitored. It was observed that, in simulation, the tumor cells tended to the capillaries as shown Figure 6 which is in line with *in-vitro* and *in-vivo* studies [26–29].

**Figure 5.**
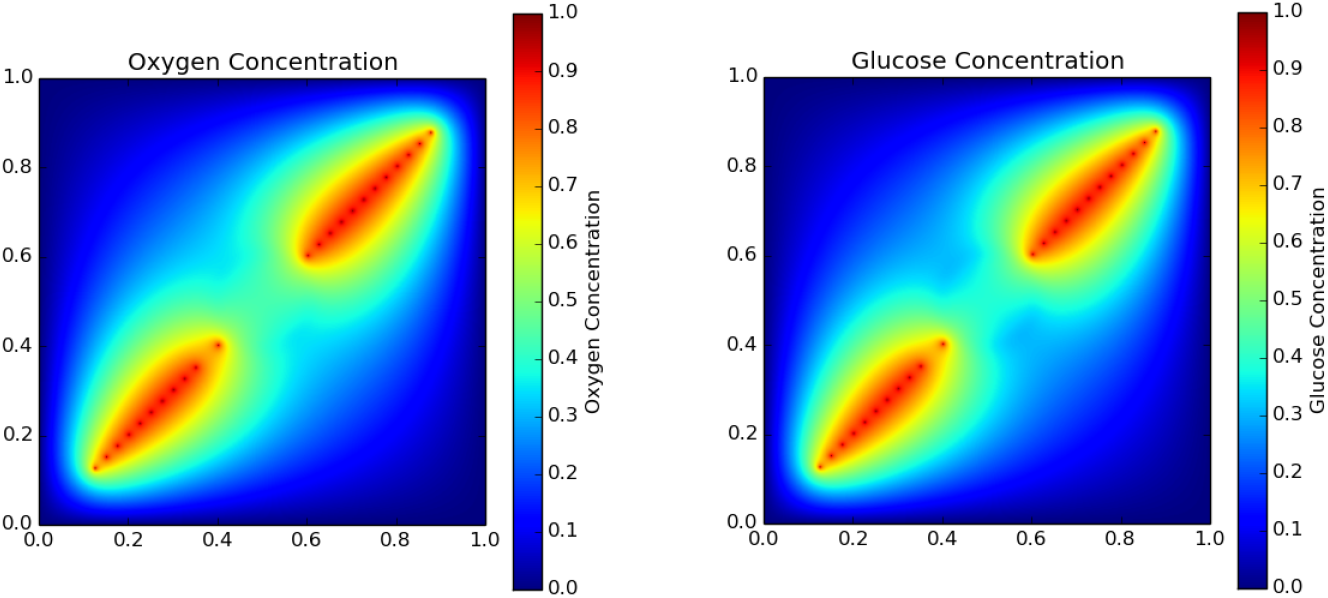
Distribution of capillaries along a diagonal line with diffusion of oxygen and glucose shown.

**Figure 6.**
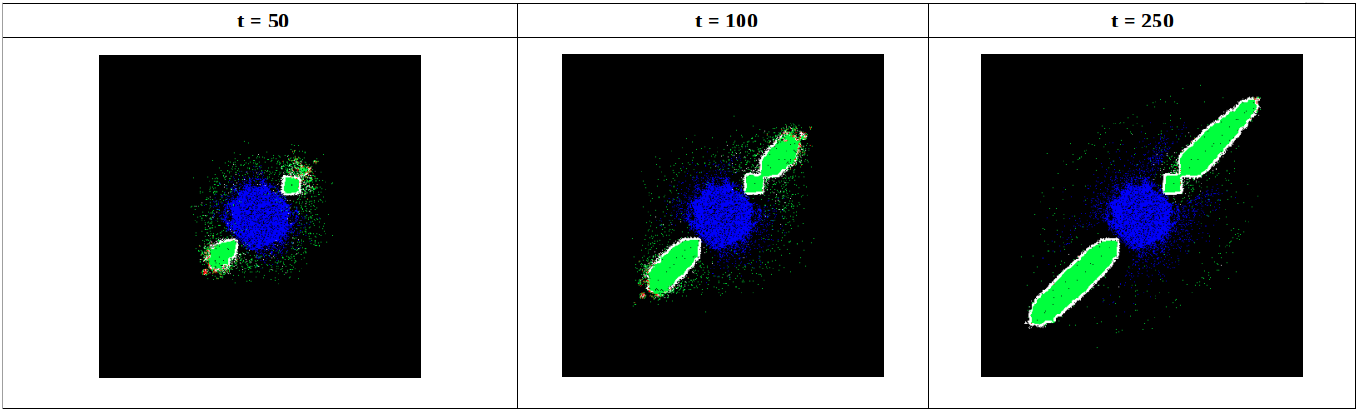
Tumor growth at time steps 5, 50 and 100 on the same diagonal axis with capillaries. Time step t equals to 22 hours.

Tumor growth patterns fit a Gompertz curve, which is a sigmoid function [30–35]. Our simulations are also validated by the observation that the tumor growth process can be defined as a sigmoid curve that starts with a high exponential growth rate but eventually levels-off with saturation (see Figure 7). Pearson correlation coefficient calculated between model and study for long term tumor growth. A strong correlation (r=0.8) for long term tumor growth is observed between tumor cell number and maximum diameter of tumor.

**Figure 7.**
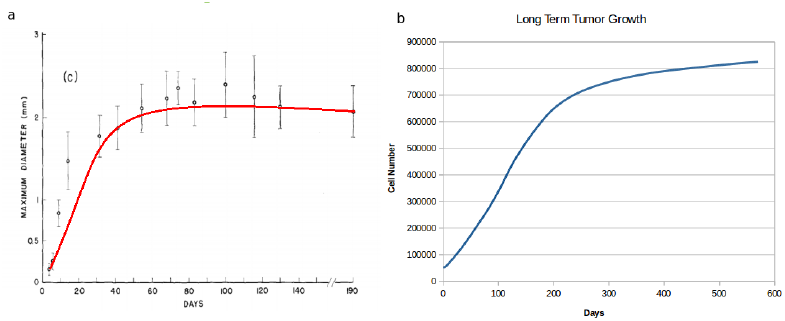
Tumor growth function fits to a Gompertzian model. **a** Experimental results [37] for long term tumor growth. **b** Results of simulation from our model

Data from Figure 1 Figure 3 and Figure 7 were extracted using Web Plot Digitizer [36] and used in calculation of the Pearson correlation coefficients.

Mutations are the foundation for the genetic structure of tumors and these are also what determined the individual differences in different cancers. In the past, some of mathematical tumor models have used this fact. Some models incorporate the effects of mutation on only one locus [38–40] while some have grouped several gene mutations according to their phenotypes [41–43]. Yet, according to the best of our knowledge, none of them inspect the effects of multiple cancer driver genes and their temporal order together, in a multiscale model. We use the results of a temporally ordered inference model generated from a multi-patient study [44]. For example, in the model, occurrence of KRAS mutation is a prerequisite for TP53 mutations and TP53 mutation is a prerequisite for the occurrence of CDK2NA mutations. In our simulations, as well, we have observed that TP53 mutations only occur after KRAS mutations and only in the cells which already have accumulated KRAS mutations. Likewise, we were able to recreate other dependecies, e.g TP53 to CDK2NA. Figure 8 shows how mutations occur in a temporal and hierarchical order, indicating that our modeling strategy is successful in simulating the actual mutation timelines in tumor progression. Using this system, a multi-clonal tumor model is created, allowing for the fact that different parts of a tumor may have different progressions of mutation.

**Figure 8.**
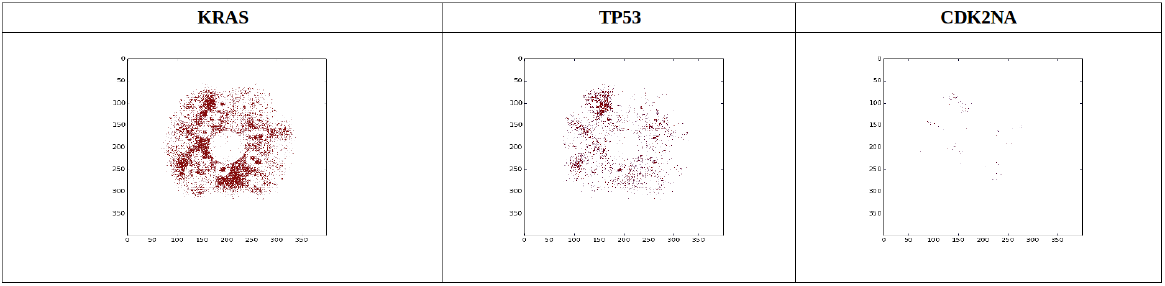
KRAS, TP53 and CDK2NA mutant cells at t=100.

### 2.2 Personalization: Estimating Parameters per Individual

Different approaches are found in the literature for the personalization of mathematical tumor models. For example, Prokopiou et. al. calculate the proliferation saturation index by using ratio of tumor volume to the carrying capacity of tumors [45]. On the other hand, Saribudak et. al. use gene expression values to personalize their model [46], and Kogan used PSA levels to individualize their model [47].

In our model, we used a stochastic decision approach in cellular automata. The decisions of each cell simulating automaton are based on the tumor microenvironment, e.g. oxygen and glucose concentrations. Weights of these variables in terms of their effect on a particular individual’s tumor are determined by using optimization. We use simulated annealing as the optimization method.

The parameter ptimization results can be seen in Figure 9. Results were obtained by optimizing parameters using the first 15 days of tumor growth data gathered from the experimental results from a xenograft [48]. Afterwards, the simulation was run with the optimized parameters. Predicted tumor growth after optimization is also shown Figure 9. The model’s prediction closely mirrors the growth trend of the xenograft.

**Figure 9.**
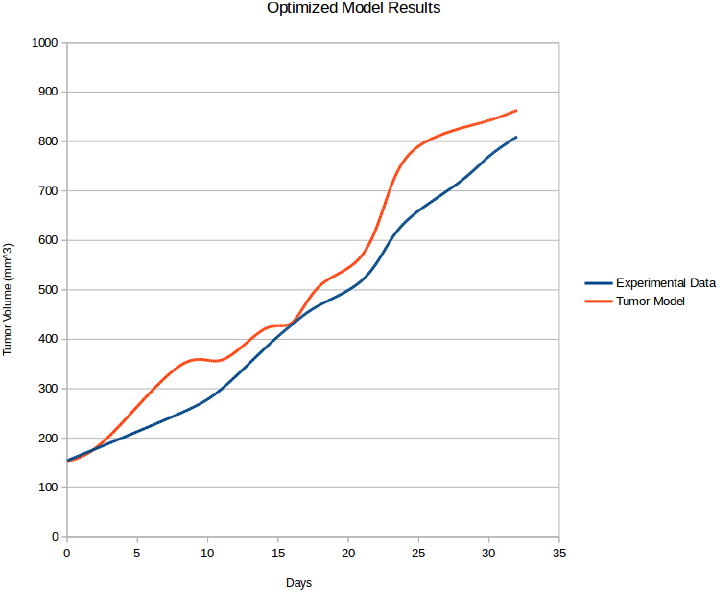
Results of a tumor growth experiment [48] and simulation results.

## Materials and Methods

### Details of the Tumor Model

We utilize the model proposed by Gerlee and Anderson [49] as a basis to our model. The Figure 10 shows the components an comprehensive tumor model should have. model.

**Figure 10.**
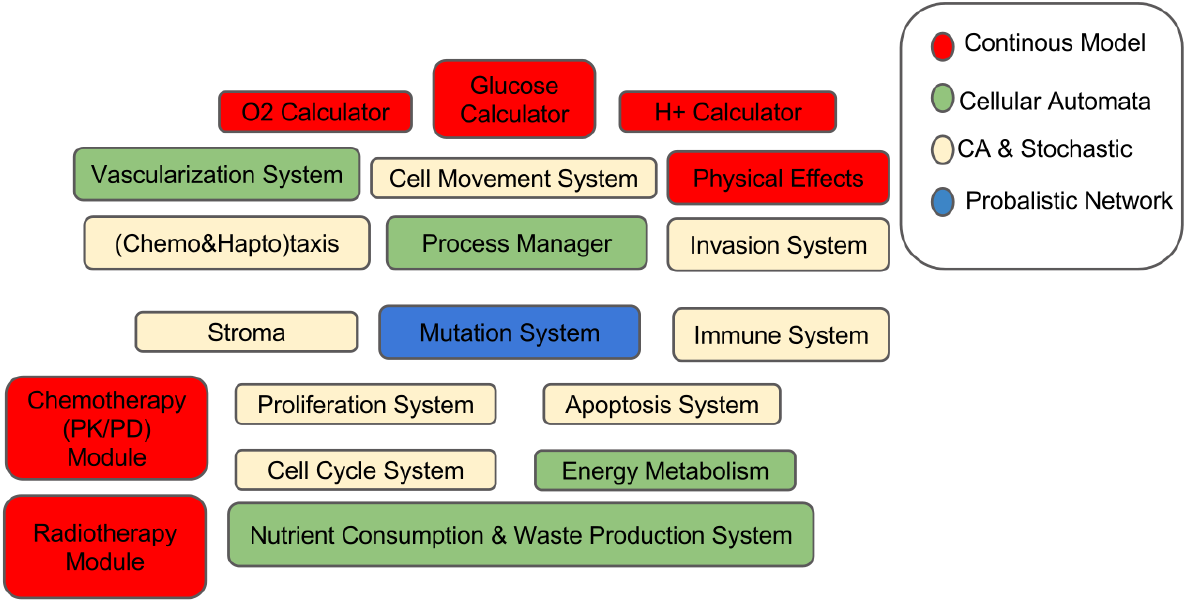
Example for a General Tumor Model

Our model is specific to lung cancer adenocarcinoma. The fixed parameters used in the model are gathered from in-vitro and in-vivo experiments from literature. The modules implemented in this study are shown in Figure 11.

**Figure 11.**
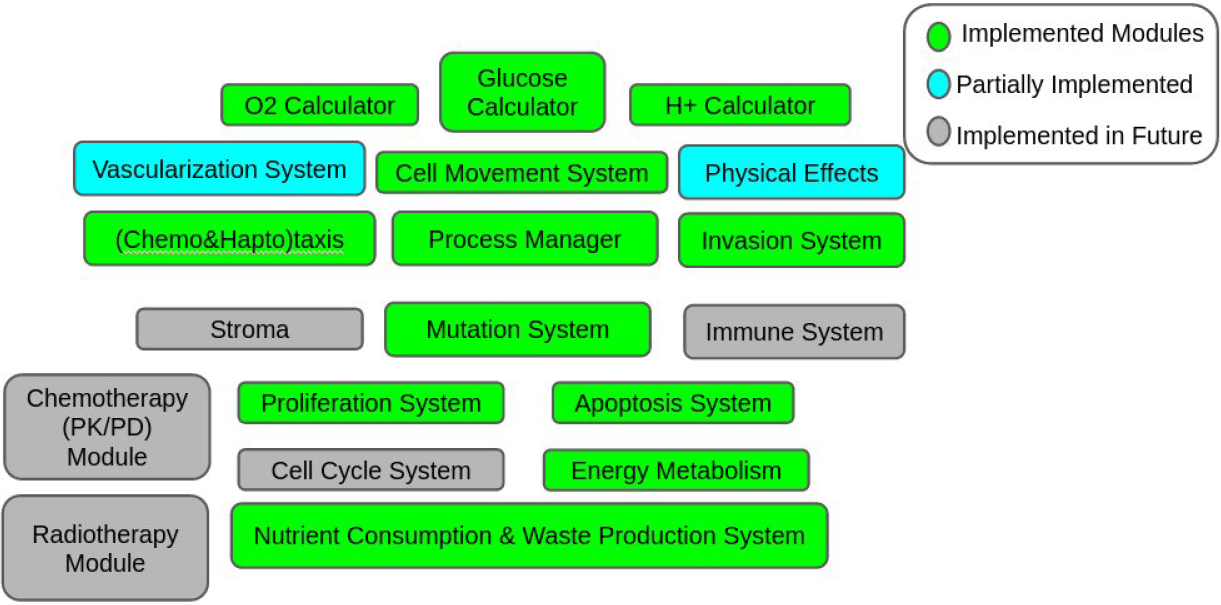
Implemented Modules in Study

Cellular automata are used to make decisions based on substance distribution and mutation effects, since rule based automata are well suited to simulating a system which depends on many variables. Decisions for migration, proliferation, apoptosis and substance consumption, are based on the state of each cellular automaton, which use a stochastic decision making process.

Each module will be discussed in the subsequent sections. In terms of the software infrastucture; Python [50] is used as the main platform for implementation of model, FiPy [51] is used to solve PDEs and Cython [52] for increased efficiency of simulation. Finally, PyGame [53] was used for implemeting the graphical user interface and for visualization.

## 3 Model Parameters

Most of the physical parameters used in our simulator are specific to lung cancer adenocarcinoma, while some are general parameters for tumors or tumor microenvironments. The A549 cell line was selected as a basis, since it represents a very common lung tumor presentation. Table 1, below, shows the parameters, symbols, and values used, with references.

**Table 1.**
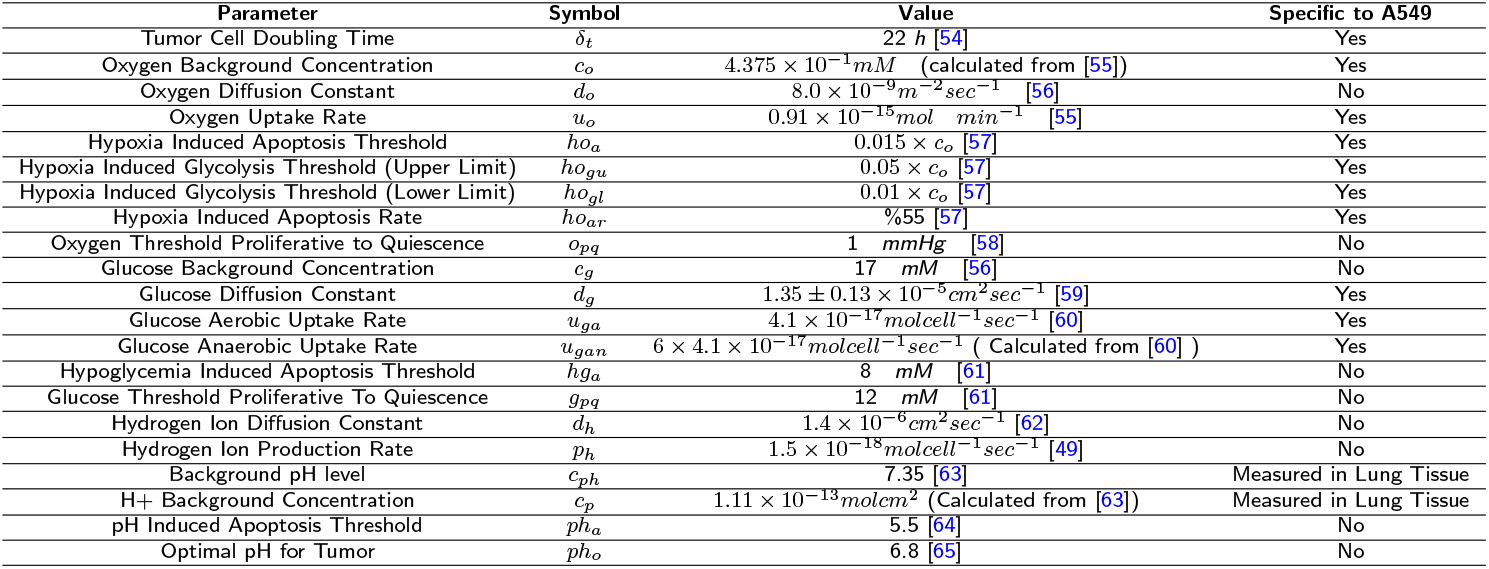
Parameters Used In Model

## 4 Diffusion of Substances in the Tumor Microenvironment

Tumor growth consists of the *consumption, growth, production, migration, apoptosis* and *necrosis* phases. Tumors need oxygen and glucose to grow and they also produce waste. If a tumor can’t find enough oxygen it goes into hypoxia. If the tumor can’t find enough nutrition (glucose), hypoglycemia starts. Producing waste (*H*+ ions) is another function of cell metabolism but also helps tumor to create an advantageous microenvironment for itself, as low pH is favored by most tumors, giving them a competitive edge over normal cells.

**Figure 12.**
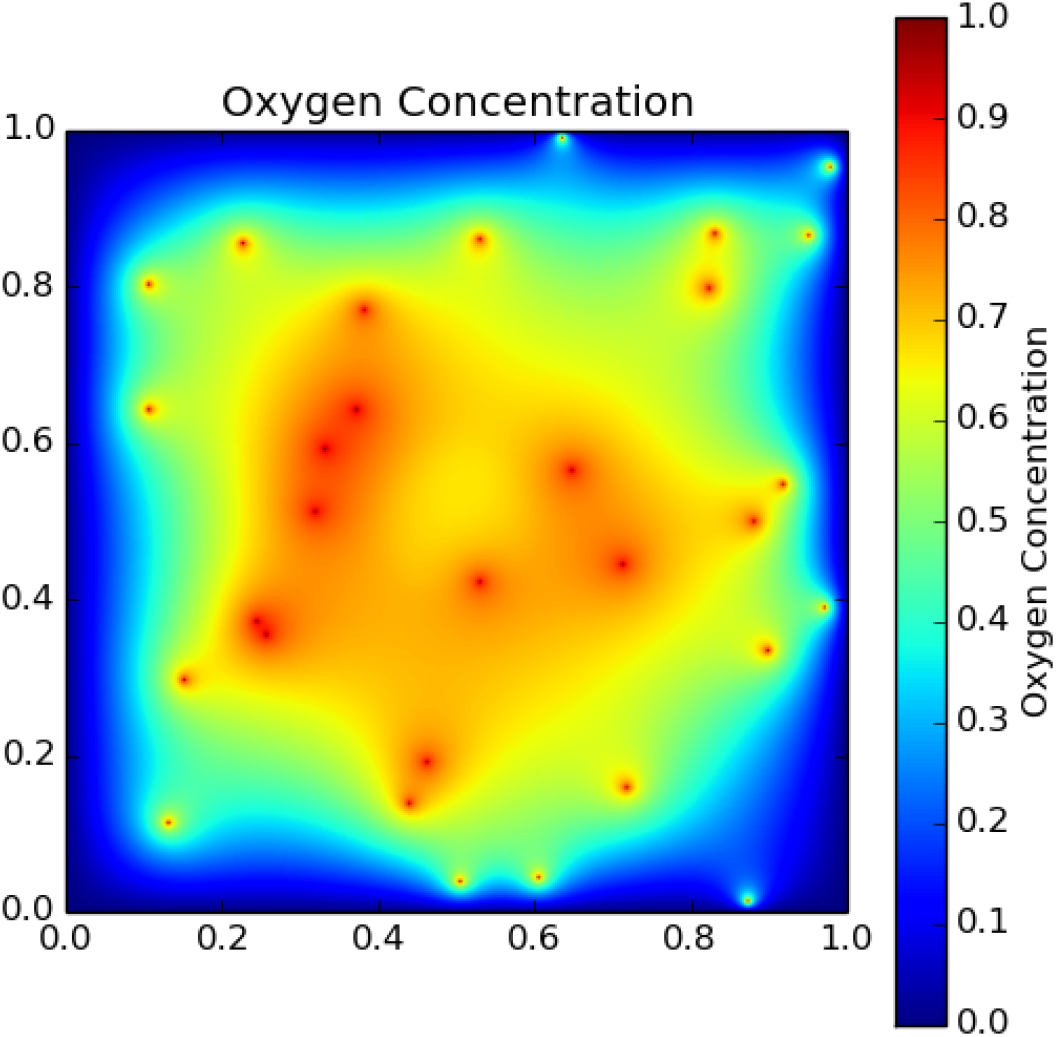
An example of oxygen distribution in cell microenvironment. Transverse sections of randomly placed capillaries are shown as red dots.

Substance diffusion into the tissue is modeled as:

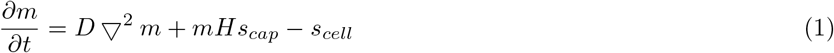

Equation 1 calculates the substance diffusion within the tissue boundaries. We assume that there are capillaries spread in tissue which supply fixed substance flow to the tissue and tumor cells which consume said substance. This equation is valid for consumed substances such as oxygen and glucose in our model. Waste substances such as H+ ions produced by tumor cells (and normal cells, alike) are, likewise, removed by capillaries. Accordingly, for waste, we modify our diffusion model as shown below to simulate this phenomenon:

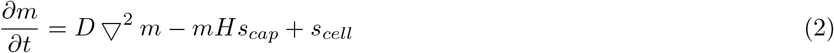

In equations 1,2 *s*_*cell*_ stands for consumption or production of substance by tumor cells and *s*_*cap*_ is the fixed term for the substance delivery or removal capacity of capillaries. Finally, H is a very large number which is used to remove effects of neighbors for the specific coordinate where the capillary exits. The particulars are elaborated on in Appendix A.

Once, the conditions of tumors’ microenvironments are thus modeled, internal dynamics of tumor cells can be explored.

## 5 Intracellular Model

Throughout their lifecycles, cells make several critical biochemical decisions. The first among these is about proliferation. Tumor cells decide weather they will proliferate, or not, based on genetic and microenvironment conditions. Another critical decision is to trigger apoptosis, if necessary. Furthermore, tumor cells manage their energy production strategy: When oxygen concentration is below a specific threshold, they change their energy metabolism from aerobic to anaerobic.

During the simulation, at each time step, each simulated cell should decide what its biochemical state is. The cell should decide on many dimensions; such as whether it will die or not, how it will produce energy, its substance consumption budget. Moreover, whether if any mutations will occur. This should be completed before other decisions are given, since nearly all other subsystems are effected by genetic variations.

After genetic changes occur, the cell starts to explore its environment to find if there is any other cells in the neighborhood and detect amount of vital substances to use them in decision making process. Then the cell decides, how it will produce energy, by using aerobic or anaerobic energy metabolisms. Finally the cell decides for its phenotype between, apoptosis, quiescence, proliferative or migrative states. Decision process is based on probabilistic functions formed by us. Weights of the variables are found using optimization which are explained in results section. Details of functions used in subsystems are explained in the following subsections. Overview of the model can be seen in Figure 13

**Figure 13.**
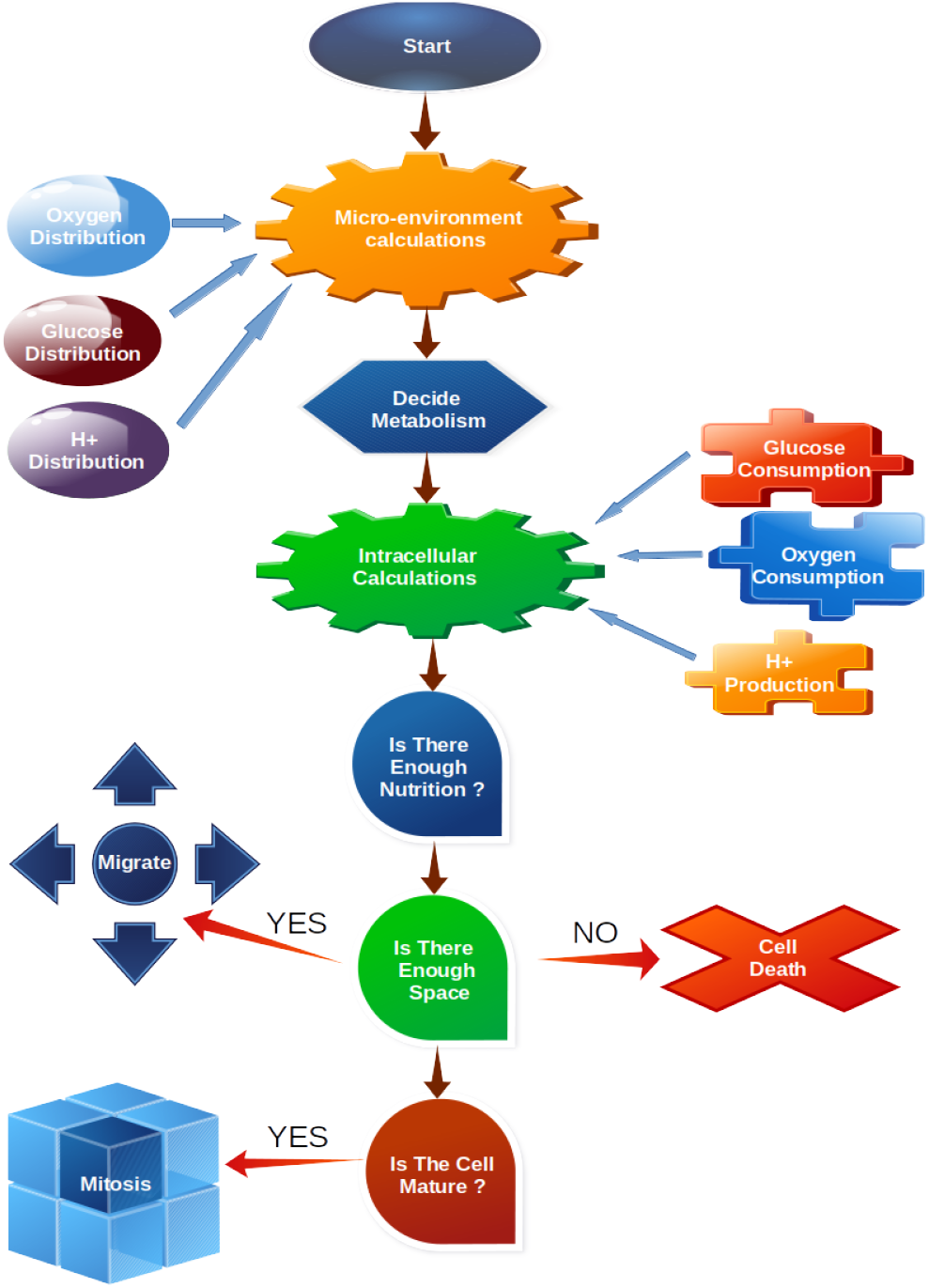
Overview of The Model

### 5.1 Proliferation

Each simulated cell’s proliferation subsystem decides whether the cell is in a proliferating state or not. When the cell is *quiescent*, if there is free space for growth, if parameters such as nutrition and oxygen level are favorable, then cell has a good chance of proliferating.

It is assumed that the maximum rate of proliferation will take place at optimal conditions. When cell’s microenvironment is closer to optimal conditions the proliferation probability increases; otherwise it decreases. Inputs used for the proliferation decision are oxygen concentration, glucose concentration and pH. Genetic effects are also used in the probability distribution function. The equation 3 is used to decide the proliferation state of the cell:

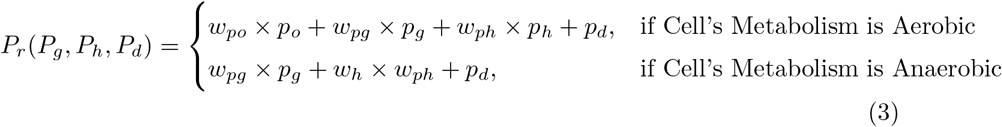

In equation 3, *P*_*r*_ represents the weighted proliferation probability, *P*_*o*_ is the probability coefficient for oxygen. *P*_*o*_ is calculated based on oxygen concentration at the coordinates of cell which represented with *m*_*o*_ and given by:

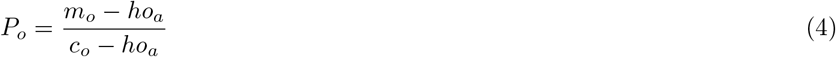

In equation 4, *ho*_*a*_ is hypoxia induced apoptosis threshold which is the minimum oxygen concentration for proliferation and *c*_*o*_ is background oxygen concentration which is the maximum oxygen concentration for cell’s microenvironment as given in Table 1. Probability coefficient for glucose *P*_*g*_, represents glucose concentration at the coordinates of the cell:

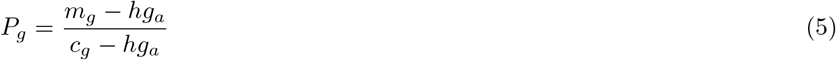

Calculation of the probability coefficient for pH, represented by *P*_*h*_, is more complex than *P*_*o*_ and *P*_*g*_, because for for oxygen and glucose higher concentration correlates with higher probability for proliferation, but for pH, the cell needs an optimal pH level to have the highest chance of proliferation. Thus, *P*_*h*_ is calculated as a piecewise function:

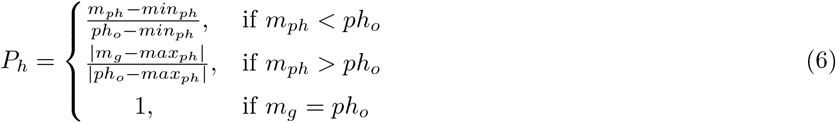

where *min*_*ph*_, *max*_*ph*_ are minimum and maximum values of pH that a tumor cell can live under; *ph*_*o*_ is the optimal pH level for proliferation as given in table 1 and *m*_*ph*_ is pH level at the coordinates of cell.

Finally, *P*_*d*_ represents effects of the cell’s accumulation of mutations on the proliferation probability and will be explained in subsection 5.8.

### 5.2 Invasion

When a cell decides on proliferation or migration, the next question is about finding the most convenient place to do so. The *invasion system* models this decisison based on microenvironment conditions. The invasion system uses oxygen, glucose and H+ concentration as input parameters. When scanning neighbour cells with traditional methods for invasion, a strange effect is occurs, as mentioned by Gerlee and Anderson [49]. The tumor tends to grow in a tree-like way, sprouting branches. Although we could not explain this effect, we overcome this issue by scanning cells orthogonally and diagonally at consequtive time steps, in interleaved fashion, as explained in Gerlee and Anderson.

To find optimal invasion coordinates, the cell prefers the direction where oxygen and glucose concentration is maximum. For H+ concentration the cell should look for optimal pH level or nearest level to optimal, given as *ph*_*o*_ in table 1. The invasion propensity score *s*_*inv*_ can then be calculated as:

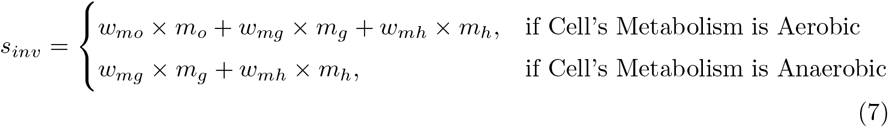

An error margin (*e*_*inv*_) should be determined which simulates the transient insensitivity of cells to proliferative oppurttunities. When maximum *s*_*inv*_ is determined, each cell’s *s*_*inv*_ is compared to the maximum *s*_*inv*_. If the difference between them is not more than *e*_*inv*_ then this cell is a candidate for invasion. After all candidates are determined, one candidate is selected randomly and a new tumor cell appears at coordinates of the chosen cell.

### 5.3 Migration

Cell migration is an important factor in the morphology of the tumor. Tumor cells may decide to migrate if conditions are not suitable to survive or proliferate. In the migration subsystem of our model, each simulated tumor cell decides whether to manifest a migrative phenotype or not. Then, it finds a suitable place for invasion by using invasion subsystem and finally it invades the location directly or by using the cell movement subsystem. The migration subsystem decides to manifest the most likely phenotype with a naive Bayes approach similar to the proliferation subsystem. There are three types of environmental parameters that force a cell to migrate. The first one is the oxygen level. When oxygen level decreases below *ho*_*a*_ then the cell’s possibility of choosing a migrative phenotype is calculated with equation 8

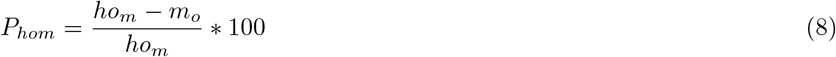

where *P*_*hom*_ stands for hypoxia based migration probability in percent, *ho*_*m*_ is hypoxia induced migration threshold which is determined based on simulation results as 5 x *ho*_*gl*_ (see table 1) and *m*_*o*_ is oxygen concentration at the cell’s coordinates. In a similar way migration probability based on glucose level can be calculated with 9:

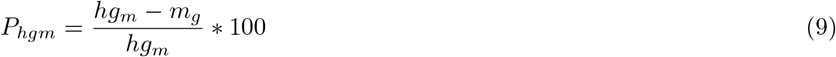

where *P*_*hgm*_ stands for hypoglicemia based migration probability in percent, *hg*_*m*_ is hypoglicemia induced migration threshold which is determined based on simulation results as 5 x *hg*_*a*_ (see table 1) and *m*_*g*_ is glucose concentration at the cell’s coordinates.

### 5.4 Apoptosis

Beyond natural apoptosis, three cases are considered for apoptosis in our model: Hypoxia, hypoglycemia and extremely low pH level can cause apoptosis.

When the oxygen level decreases below %1.5, hypoxia starts until oxygen runs out (%0). It has been shown that %55 of NSCLC Adenocarcinoma (A549) cells die when oxygen level reaches %0. Also, it is known that natural apoptosis rate is %10 for A549 cells [57].

Since 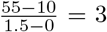 for each %0.1 change at oxygen level, apoptosis survival probability increases %3. Based on this assumption, hypoxia based apoptosis probability can be calculated as:

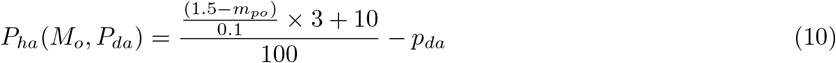

where oxygen concentration at the coordinates of the cell are represented with *m*_*po*_ and probability of as genetic effects decreases apoptosis chance and is represented by *p*_*da*_.

Cells can live without oxygen but not without glucose. Hypoglycemia induced apoptosis probability (*P*_*ha*_) can be calculated when *m*_*g*_ < *hg*_*a*_ as follows:

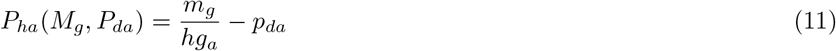

The last factor which causes apoptosis is pH level. An acidic microenvironment is favorable for the tumor because cancerous cells are more resistant to acidic environment than parenchyma. But, when pH level decreases to severly low levels all kinds of cells starts to die. When this effect is modeled, pH level at the coordinates of the cell is represented by *m*_*h*_ and *min*_*ph*_ stands for the minimum pH level of cell microenvironment. Thus, we can calculate *P*_*ha*_ (apoptosis probability based on pH) as:

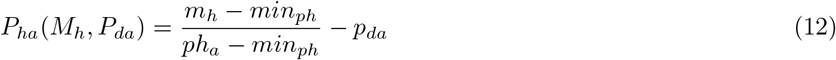

Apoptosis probability is calculated using oxygen, glucose and pH level. However, even as the cell decides to start apoptosis it will wait for a period. This delay act as a low-pass filter and prevents the cell from being affected by noise and momentary oscillations of microenvironment signals. There are more complicated methods that can be found in literature to model delayed systems [66].

### 5.5 Energy Metabolism

Tumor cells can produce energy by using two different metabolisms; aerobic and anaerobic. In aerobic metabolism, oxygen and glucose are used to produce energy:

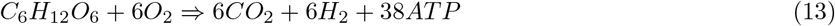

In the anaerobic metabolism, only glucose is used to produce energy:

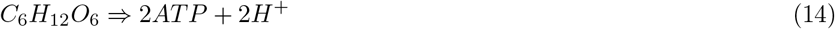

In our model, the energy metabolism shift is only based on oxygen concentration; genetic effects are ignored for the sake of simplicity.

Metabolism shift starts at %5 oxygen concentration with %20 probability and reaches %100 probability at %1 oxygen concentration [57]. If the concentration is over %5 then cell always chooses aerobic metabolism. Oxygen concentration at coordinates of cell as percentage (*m*_*po*_) is obtained from:

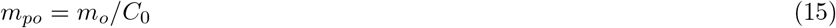

and the probability of a cell’s metabolism change from aerobic to anaerobic is calculated by:

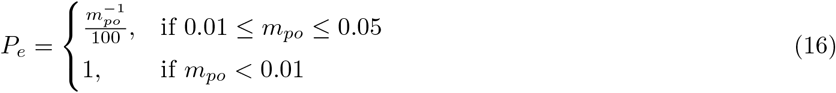

There is also a delay introduced before the decision for a metabolism shift, with the same rationale as for the delay for the apoptosis decision, explained above.

### 5.6 Oxygen - Glucose Consumption and Acid Production

Tumor cells have a baseline oxygen consumption rate, represented by *u*_*o*_ as seen in Table 1, but this rate changes based on cellular conditions. For example a cell will not use oxygen when under anaerobic metabolism. Also, the cell’s state will effect oxygen consumption. We assume that the cell consumes %50 more oxygen in proliferative state than in the quiescent state.

Glucose consumption (*u*_*g*_) is a calculated in a similar way. Glucose consumption in aerobic and anaerobic states are calculated based on the stochiometry of the respective metabolisms. Proliferative cells are assumed to use %50 more glucose, similar to oxygen.

Acid production is observed in the anaerobic metabolism, due to glycolysis. Hydronium production rate (*p*_*h*_) is taken from literature [49]. At proliferative state, hydronium production is assumed to increase by %50, since glucose consumption increases with same ratio.

### 5.7 Cell Stress and Movement

Physical simulations on tumors shows us that there is a proliferative belt on tumor mass. This belt is caused due to various reasons, e.g. cell-cell adhesion and cell-ECM adhesion [4]. In our model, instead of treating each physical effect one by one in detail, we use a *stress score* variable.

The stress score for each cell is calculated proportional to the cell’s distance from tumor’s edge. Pressure on cells located in deeper parts of tumor will be higher than cells which are located at the edge of the tumor because of cell-cell adhesion and cell-ECM adhesion. Edge cells’ tendency of migration is thus higher than deep cells.

### 5.8 Genetic Effects on Tumor

When modeling cancer, simulating genetic effects, taking in to account all variations is intractable. Since the aim of this study is only creating a simple proof of concept model, just a few important mutations of A549 are included, based on literature [44].

Our model uses an inheritance mechanism. When a simulated cell proliferates, it copies the DNA of its ancestor, so mutations are transferred between generations. As a first step, relationship between mutations determined. After this step, probabilities of mutations calculated. Finally, effects of the mutations are added to the model. In the model, only two types of mutation effects are considered: mutations’ effects on probabilities of proliferation and apoptosis.

For both, if any mutation occurs in a cell, each driver mutation provides only a small selective growth advantage to the cell, on the order of a 0.4% increase in the difference between cell birth and cell death [67]. Since only 5%-10% of these are driver mutations, we calculate each driver will have an effect of 8%. Thus, if any mutation occurs in a cell, it is assumed that proliferation probability increases 8% or apoptosis probability decreases 8% based on mutation type. These relationships and effects are shown in Table 2 with minimum and maximum probabilities of observing said mutations in the population.

**Table 2.** Mutations of A549 Used In Model

| Mutation | Minimum Probability | Maximum Probability | Ancestors | Effect Type |
| --- | --- | --- | --- | --- |
| EGFR [68] | 10 | 35 | None | Proliferation |
| KRAS [69] | 33 | - | None | Proliferation |
| NTRK3 [70] | 3 | - | None | Proliferation |
| TP53 [70] | 30 | 50 | KRAS | Apoptosis |
| ATM [70] | 10 | 20 | KRAS | Proliferation |
| STK11 [70] | 18 | - | TP53, NTRK3 | Apoptosis |
| CDK2NA [70] | 5 | - | TP53 | Apoptosis |

The model for the mutations is based on a Bayesian network. At each time step, a simulated cell triggers its own mutation system. For each mutation, preconditions are checked. For example for A549, EGFR mutation almost never occurs in tumors with KRAS mutation or TP53 mutation occurs if KRAS gene has a mutation. For the sake of the simplicity we assume that these mutations are all pathogenic.

After ancestors of the mutation are validated, a mutation probability is determined for each viable mutation. Finally, if a mutation occurs, effects of said mutations, whether on proliferation or apoptosis, are applied to the cell in quesiton, and will affect its future decision equations.

## 6 Conclusion and Future Work

Literature includes many studies on personalized medicine, but most of these are results of bulk biostatistics and bioinformatics analyses. Tumor models on the other hand, have potential in providing mechanistic and simulated personalized predicitons. In this study, we demonstrate a tumor model based on empricial observations and biochemical properties from previous litarture that is amenable to tuning and extension, and able to provide personalized predictions.

To the best of our knowledge, our model is the first one that can predict personalized growth patterns of lung adenocarcinoma and validate it with xenograft model data. A hierarchical Bayesian network modeling the genotypes of tumor subpopulations was also created to predict ordering and timing of mutations during tumor progression.

We validated our model’s behavior with experimental data from literature. We observed that both under hypoxia and hypoglicemia our model’s growth pattern is in line with experimental results. Furthermore, our model shows the expected Gompertzian curve in chemotactic behaviour and in general growth pattern. Our personalization strategy is also promising. We have shown that if a model consistent with tumor biology can be developed, it can be personalized using a simple parameter optimization method like simulated annealing.

In the future we will extend our model by simulating effects of the immune system and integrate chemotherapy, radiotherapy and immunotherapy results to our model towards developing a clinical decision support tool.

## Appendix A

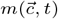 means concentration of substance at coordinates 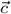 at time *t. D* is diffusion constant of substance. Since our model is 2D,

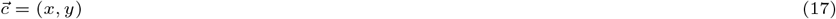

we could know concentration of substance at specified coordinates at time *t*. To solve equation it could also be written as below:

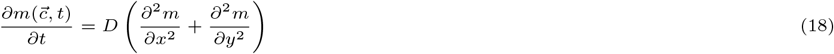

A detailed explanation for this equation could be found in literature [71]. Physical respresentation of diffusion phenomenon which is respresented as equation 18 could be seen in figure:

**Figure 14.**
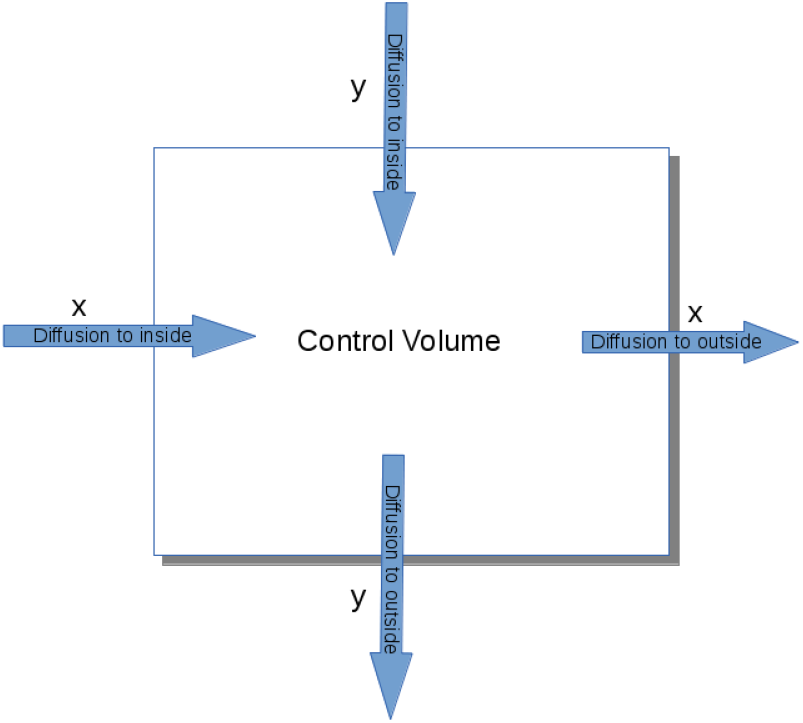
Diffusion as a Physical Phenomenon

After the basic equation of diffusion is formed, only a source term is needed which represents consumption or production of the substance for a coordinate at a specific time [71]:

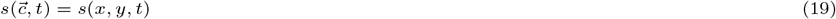

Oxygen and glucose which are consumed by cells are subtracted from equation 18:

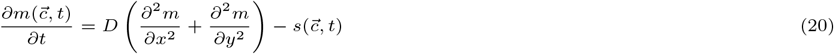

and source term for hydrogen ions (**H**+) which are produced at the end of glycolysis is added:

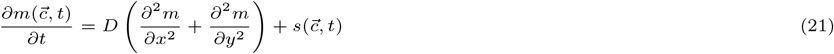

Now effects of capillaries should be added to equation. We accomplish this by using a implicit source term which is a product of a fixed value with diffusion term (m). We also multiply it with a huge value (**H**) to winnow effects of neighbors out. We write the final equation for consumed substances as:

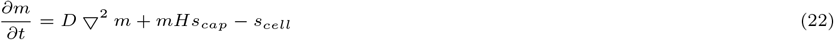

for produced substances it will be:

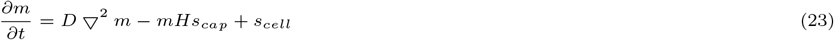

Finally, initial and boundary conditions for equations 20 - 21 will be determined [72]. We assume that boundaries of grid with *L × L* size, concentration equals to 0. Initial conditions could be defined as

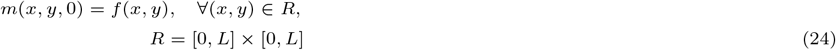

with Dirichlet boundary conditions

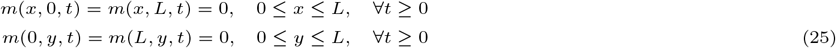

### 6.1 Non-dimensionalization

Non-dimensionalization can be done with dividing each term of an homogeneous equation to parameters which have same units [73]. In this study, diffusion constants and consumption/production rates were non-dimensionalized with the following equations:

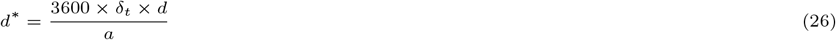

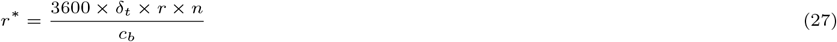

These are generalized form of non-dimensionalization equations. In equation 26, *d*\* is non-dimensional diffusion constant. Tumor cell doubling time (*δ*_*t*_) multiplied by 3600 to convert hour to second. Original diffusion constant is represented with *d*. Area (*a*) is total area (in *cm*^2^) of the grid. In equation 27, *r*\* represents non-dimensional consumption/production rate. Original rate is represented by *r* Maximum number of tumor cells (which equals to number of grid’s cells) is represented by *n*. Finally background concentrations are represented by *c*_*b*_. Using these equations 26 - 27, *u*_*o*_, *u*_*ga*_, *u*_*gan*_, *p*_*h*_, *d*_*o*_, *d*_*g*_, *d*_*h*_ were non-dimensionalized.

## Code Availability

Source code is available at: https://gitlab.com/serbulentu/MultiscaleTumorModel

